# Tissue interactions govern pattern formation in the posterior lateral line of medaka

**DOI:** 10.1101/2020.03.26.009969

**Authors:** Ali Seleit, Karen Gross, Jasmin Onistschenko, Oi Pui Hoang, Jonas Theelke, Lázaro Centanin

## Abstract

Vertebrate organs are arranged in a stereotypic, species-specific position along the animal body plan. Substantial morphological variation exists between related species, especially so in the vastly diversified teleost clade. It is still unclear how tissues, organs and systems can accommodate such diverse scaffolds. Here, we use the sequential formation of neuromasts in the posterior lateral line (pLL) system of medaka fish to address tissue-interactions defining a pattern. We show that the pLL pattern is established independently of its neuronal wiring, and demonstrate that the neuromast precursors that constitute the pLL behave as autonomous units during pattern construction. We uncover the necessity of epithelial integrity for correct pLL patterning by disrupting *keratin 15* (*krt15*) and creating epithelial lesions that lead to novel neuromast positioning. By using *krt15/wt* chimeras, we determined that the new pLL pattern depends exclusively on the mutant epithelium, which instructs *wt* neuromast to locate ectopically. Inducing epithelial lesions by 2-photon laser ablation during pLL morphogenesis phenocopies *krt15* genetic mutants and reveals that epithelial integrity defines the final position of the embryonic pLL neuromasts. Our results show that a fine-balance between primordium intrinsic properties and instructive interactions with the surrounding tissues is necessary to achieve proper organ morphogenesis and patterning. We speculate that this logic likely facilitates the accommodation of sensory modules to changing and diverse body plans.

## Introduction

Among the members of an animal species, organs are arranged in fixed numbers and stereotypic positions along the body. Subtle or major variations that occur between related species have necessarily evolved by maintaining organ functionality and association with the rest of the tissues in the organism. There is a plethora of body sizes and patterns in the animal kingdom, where teleost fish in particular have mastered body alteration and re-shaping. How the different tissues, organs and systems can scale and accommodate to such diverse scaffolds is still poorly understood. So is the impact of changes in one tissue on another, *i.e* the hierarchical organisation governing these changes.

The reproducibility of developmental programs during organogenesis appears to be an intrinsic property of the system, as self-organising programs were revealed using different organoid models during the last decade (Eiraku et al., 2011; Lancaster et al., 2013; Sato et al., 2009; Turner et al., 2016; van den Brink et al., 2014). Organ location, on the other hand, is achieved in two different manners depending on the animal model. Cases like *C. elegans* constitute a clear example of a fixed, deterministic lineage where the precise temporal and spatial order guarantees the formation of cell types and tissues in a *defined* location (Sulston et al., 1983). Most vertebrates analysed follow another developmental logic, though, where an initial symmetry break will induce the appearance of different cell types, tissues and eventually organs (Martinez Arias and Steventon, 2018). Induction events constitute examples of how cell types and tissues interact with each other to instruct their timely appearance at *relative* locations. The rationale that emerges for vertebrate organogenesis is that of an initial plasticity - *where* the tissue will be formed - followed by a fixed, self-organising programme - *how* the organ will form. How variations in tissue interactions could result in new patterns during development remains largely unknown.

The lateral line is a sensory system whose organs, the neuromasts, distribute along the surface of fish and amphibia (Sapede et al.,2002; Ghysen and Dambly-Chaudière, 2007). Their exposed location and unique morphology have made them a popular model to tackle several aspects of organogenesis. Many groups have contributed to our extensive understanding of the signalling pathways shaping the embryonic lateral line system in zebrafish (Aman and Piotrowski, 2008; Chitnis et al., 2012; David et al., 2002; Grant et al., 2005; Haas and Gilmour, 2006; Hernández et al., 2006; Lecaudey et al., 2008; López-Schier and Hudspeth, 2005; Lush and Piotrowski, 2014; Ma et al., 2008; Nechiporuk and Raible, 2008; Pinto-Teixeira et al., 2015; Romero-Carvajal et al., 2015; Sánchez et al., 2016; Wada et al., 2013b; Wibowo et al., 2011). Briefly, a primordium migrates concurrently with the pLL nerve and associated glia along the horizontal myoseptum and deposits a handful of pre-formed neuromasts from its rear edge (Ghysen and Dambly-Chaudière, 2007; Grant et al., 2005; Lecaudey et al., 2008; López-Schier and Hudspeth, 2005; Lush and Piotrowski, 2014; Nechiporuk and Raible, 2008). Additional primordia will migrate later, expanding the initial set of organs during the larval stages (Ghysen and Dambly-Chaudière, 2007; Ledent, 2002; Nuñez et al., 2009; Sapède et al., 2002; Whitfield, 2005). Some aspects of lateral line formation have been reported as well in different fish species, including tuna, *Astyanax* and medaka (Ghysen et al., 2012; Ghysen et al., 2010; Pichon and Ghysen, 2004; Sapède et al., 2002; Seleit et al., 2017a; Seleit et al., 2017b). The availability of genetic tools in medaka allowed a dynamic study of neuromast generation, which revealed that one primordium is responsible for the sequential generation of two different, parallel lateral lines (Seleit et al., 2017a). A first set of neuromasts is deposited by the primordium (*primary* neuromasts) and move ventrally immediately after deposition to form the ventral pLL. Later, a second set of organs (*secondary* neuromasts) is formed between each pair of primary neuromasts and migrate dorsally to form the midline pLL (Figure 1). The posterior lateral line in medaka at the end of embryogenesis, therefore, represents a unique neuromast pattern that deviates from that of previously studied fish.

**Figure 1.**
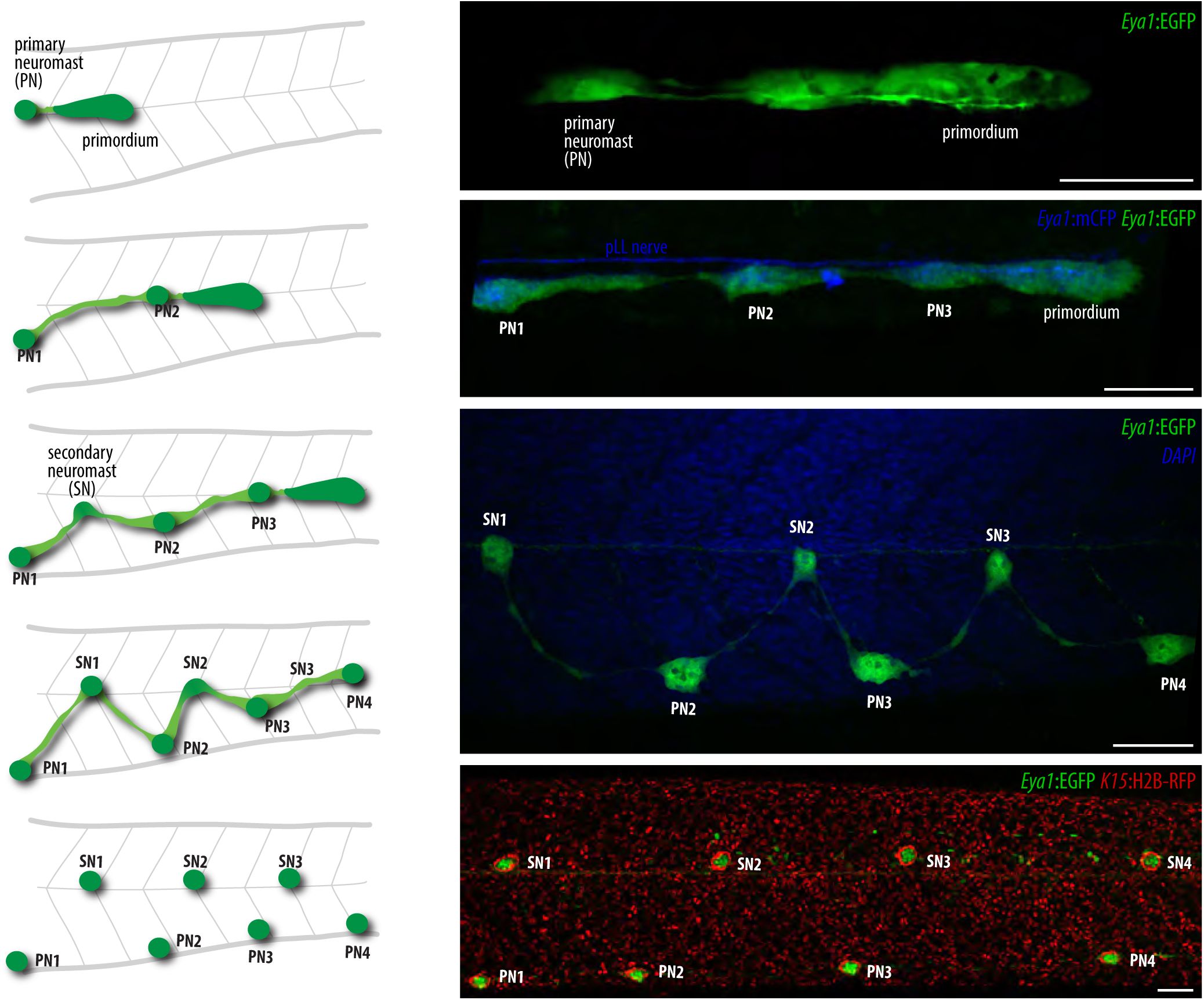
The formation of the posterior lateral line (pLL) system in medaka. A primordium (green, upper panel) detaches from the pLL placode and migrates posteriorly, with the pLL following its path (cyan, the second panel, right). Eya1 is expressed in the primordium, the deposited neuromasts and also by neurons in the ganglia. Primary neuromasts (PN) are deposited at roughly regular intervals in the midline, and move ventrally immediately after deposition (upper and second panels). When in the ventral domain, secondary neuromasts (SN) form between each pair of primary neuromasts, and these SN migrate dorsally reaching the midline to form a characteristic zig-zag pattern (third and fourth schemes, left, and third panel, right). By the end of embryogenesis, the pLL is composed of two parallel lines: a ventral pLL formed by primary neuromasts, and a midline pLL formed by secondary neuromasts. Scale bar = 50 microns. Anterior is to the left and dorsal is up in all panels.

Here, we use the sequential formation and opposite migration of neuromasts in medaka to address the tissue interactions necessary to build a pattern. By using 2-photon laser ablations we discard any role for the lateral line nerve and the associated glial cells in pattern formation and organ numbers. We then analyse transgenic lines labelling the pLL of medaka mutants with duplicated body sectors to demonstrate that primary neuromasts resolve organ positioning in an autonomous manner. Despite autonomous neuromast location, our 4D analysis reveals intrinsic properties of the pLL system. Namely, one secondary organ forms in between two primaries, and all secondary neuromasts inevitably locate to the midline, irrespective of the relative position of their founder cells. We also generate viable *keratin 15* mutants to reveal that the integrity of the epithelium influences the embryonic pattern of the medaka pLL. We confirm these results by mechanically interfering with epithelial integrity locally and recapitulate the patterning defects observed in our *krt15* mutants. Our results strongly suggest that tissue interactions govern lateral line pattern formation and that patterning in this system is resolved locally and autonomously by neuromast organ precursors. Pattern construction in the medaka pLL is governed by a balance of intrinsic primordium properties and extrinsic influences from its immediate environment, where the modulation of the latter can easily lead to novel pLL patterns. We speculate that this plasticity is exploited evolutionarily to generate novel pLL patterns accommodating the diverse sizes and body-plans of teleost fish.

## Results

### pLL nerve and associated glia dispensable for correct pLL pattern and organ numbers in medaka

The embryonic pLL pattern construction in medaka involves highly stereotypic organ movements and positioning. Primary neuromasts are deposited at the midline and will end up ventrally, while secondary organs, formed ventrally, will locate to the midline (Seleit et al., 2017a). It is still unclear what drives the ventral movement of primary organs during the initial phases of pLL pattern formation. A major difference between primary deposited organs from the primordium and secondary forming ones is that primary neuromasts are connected to the pLL nerve as they are formed (Seleit et al., 2017a). To test whether the pLL nerve can act instructively to guide the positioning of primary organs to the ventral side we decided to target the nerve by two-photon laser ablation. We utilized *Eya1*:mCFP, *K15*:H2B-RFP embryos, where membrane-tagged cyan fluorescent protein guarantees a clear labelling of the nerve, and the nuclear tagged red fluorescent protein labels the mature neuromasts of the pLL (Figure 2A-A’). We performed a high precision injury to a specific segment of the pLL nerve in 3-4 dpf embryos, when the primordium has already deposited a number of primary organs at the midline (Figure 2B-B’’). We were able to leave the primordium, the deposited neuromasts and the un-targeted anterior segment of the pLL nerve intact and uninjured (Figure 2B’’). Intriguingly, 5 days post injury there were no signs of nerve cone regrowth suggesting a deficiency in the regenerative potential and axonal guidance in the peripheral nervous system of medaka (Figure 2C). Despite the irrevocable loss of the pLL nerve neither organ numbers nor positioning were affected when fish with ablated pLL nerves were analysed at later stages of embryonic development (Figure 2C, C’ n=11 embryos). These larvae displayed the characteristic alternated pattern between primary ventral and secondary midline neuromasts. Our results indicate that the pLL nerve is dispensable for correct patterning in the pLL of medaka.

**Figure 2.**
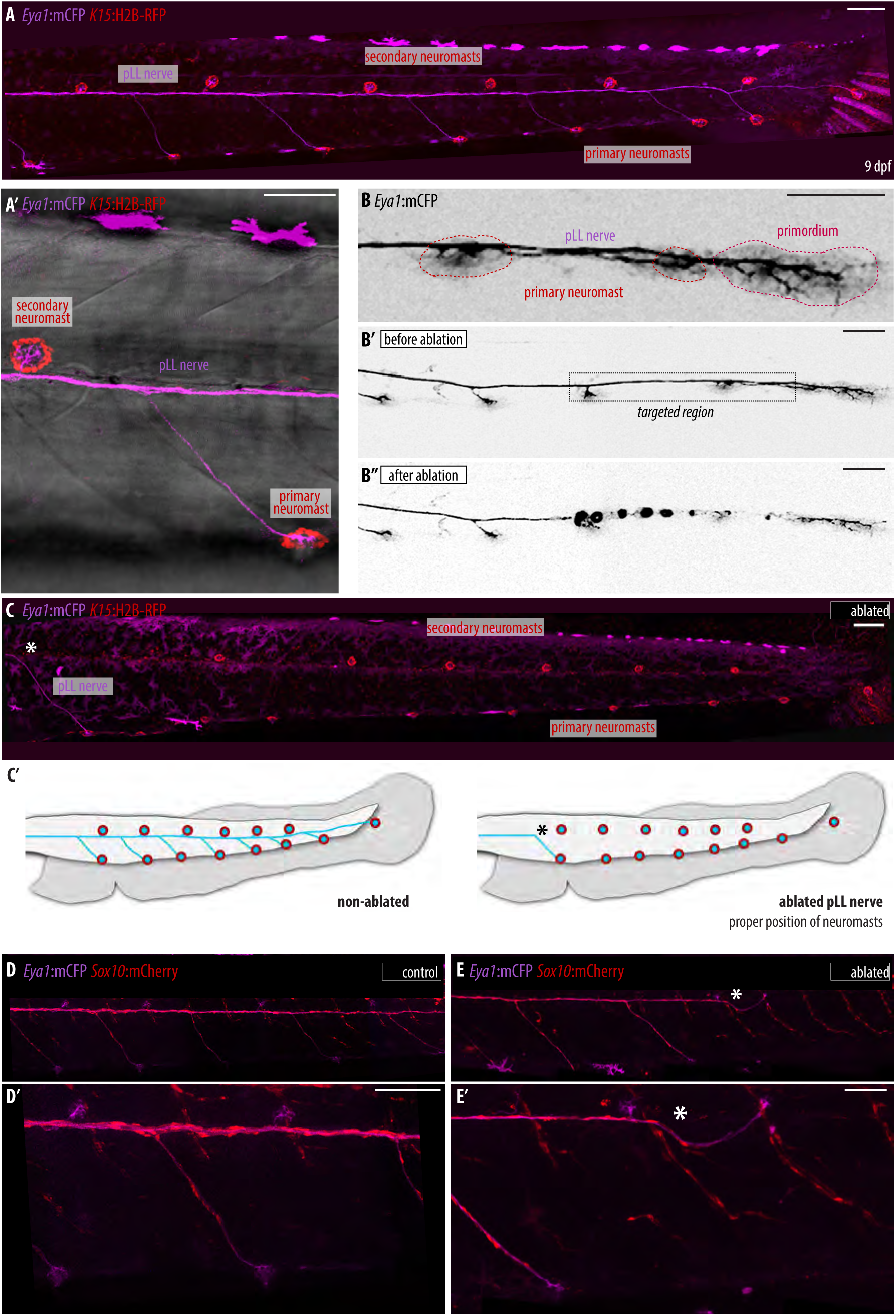
Correct organ number and positions in pLL of medaka despite loss of pLL nerve. (**A-A’**) Control 9 dpf Tg(*Eya1*:mCFP)(*K15*:H2B-RFP) embryo displays alternating neuromasts in the pLL. The pLL nerve is labelled by *Eya1*:mCFP along the trunk and is connected to the mature primary ventral organs and midline secondary organs that are both labelled by *K15*:H2B-RFP. (**B**) 3-4 dpf Tg(*Eya1*:mCFP) medaka embryo highlighting the primordium, the pLL nerve and deposited primary neuromast organs along the horizontal myoseptum. (B’) 3-4 dpf Tg(*Eya1*:mCFP) medaka embryo before multi-photon laser ablation, primordium is migrating and a few organs have been deposited. (B’’) 3-4 dpf Tg(*Eya1*:mCFP) medaka embryo after multi-photon ablation, segment of the pLL is nerve ablated while neuromasts and primordium remain intact and un-injured. (**C**) Pattern and number of neuromasts are normal despite loss of pLL nerve in Tg(*Eya1*:mCFP) (*K15*:H2B-RFP) 9 dpf. (N = 11 embryos). White asterisk indicates ablation site. (C’) Scheme of neuromast position and number upon ablation of the pLL. (**D-D’**) Double transgenic (*Eya1*:mCFP)(*Sox10*:mCherry) 9 dpf embryo shows glial cells (red) wrapping around pLL nerve (magenta). (**E-E’**) Ablated pLL nerve in Tg(*Eya1*:mCFP)(*Sox10*:mcherry). Note that the Cherry positive glial cells do not over-take the ablated pLL nerve (white asterisk indicates ablation site) and are only present where the nerve is intact (N=4 embryos). Scale bar = 100 microns.

Our results are striking since it has been extensively reported in zebrafish that loss of the pLL nerve leads to supernumerary organ formation in the lateral line (Grant et al., 2005; López-Schier and Hudspeth, 2005; Lush and Piotrowski, 2014; Whitfield, 2005). This is primarily due to the fact that associated glial cells are not able to migrate in the absence of the pLL nerve (Gilmour et al., 2002) and are thus incapable of maintaining neuromast organ numbers in check (Grant et al., 2005; López-Schier and Hudspeth, 2005; Lush and Piotrowski, 2014; Whitfield, 2005). To test whether glial cells in medaka can migrate independently of the nerve and hence rescue the possible patterning and organ number defects we generated a *Sox10*:mCherry transgenic line that labels glial cells (Lush and Piotrowski, 2014). We used double positive (*Eya1*:mCFP)(*Sox10*:mcherry) embryos to label both the pLL nerve for the ablation and the glial cells for the response after injury (Figure 2 D-E’).

As has been reported in zebrafish glial cells do not over-take the pLL nerve (Gilmour et al., 2002) and were only present on intact uninjured segments of it (N=4 embryos) (Figure 2 E-E’). These results strongly argue that the pLL pattern and organ numbers are properly established even in the absence of sox10+ glial cells and the pLL nerve in medaka.

### Uncoordinated neuromast patterning in *Da*, ventral-mirrored embryos

Primary neuromast positioning in medaka could either be a result of an active migration or a passive displacement ventrally after deposition. To distinguish these possibilities, we made use of the *Double anal* (*Da*) medaka mutant, which has a partial duplication of the ventral side along the tail due to the inactivation of *zic* genes (Moriyama et al., 2012; Ohtsuka et al., 2004) (Figure 3A-B). This is particularly interesting as it meant we could modulate the immediate environment the primordium travels through without changing any intrinsic properties of the primordium itself. If primary organs are passively displaced ventrally then we would expect a randomization of their positions in the *Da* mutant; either to the induced ventralized side or the regular ventral side. However, what we observed is that in 80% of all larvae one or more primary organs is positioned at the midline (Figure 3C-D N>50 embryos). This argues against a passive displacement of primary organs and suggests active migration (for e.g in response to a duplicated chemokine signal emanating from the ventral side) could be responsible for organ positioning. It is possible that primary organ positioning is resolved coordinately on a system level, alternatively organ positioning could be resolved autonomously by each organ (neighbor-independent movement). To distinguish whether the pLL pattern is built on a system level control or by autonomous units that undergo the same morphogenetic movements we decided to perform long term live-imaging on *Da* mutants during pLL pattern construction. This revealed that primary precursor organs are autonomous in the direction of their movement even within the same pLL and that the pLL nerve connection follows the migratory path of the forming organs irrespective of their migratory direction (Figure 3E-E’) (Supplementary Movie 1, N=5 embryos). In addition, organs that remain at the midline commonly changed their positions multiple times (above and below the horizontal myoseptum) before eventually settling at the midline (Supplementary Movie 1, 2 N = 5 embryos). Our results show that pattern construction in the pLL of medaka is highly dependent on local interactions that are resolved individually by the primary precursor organs depending on their immediate environment.

**Figure 3.**
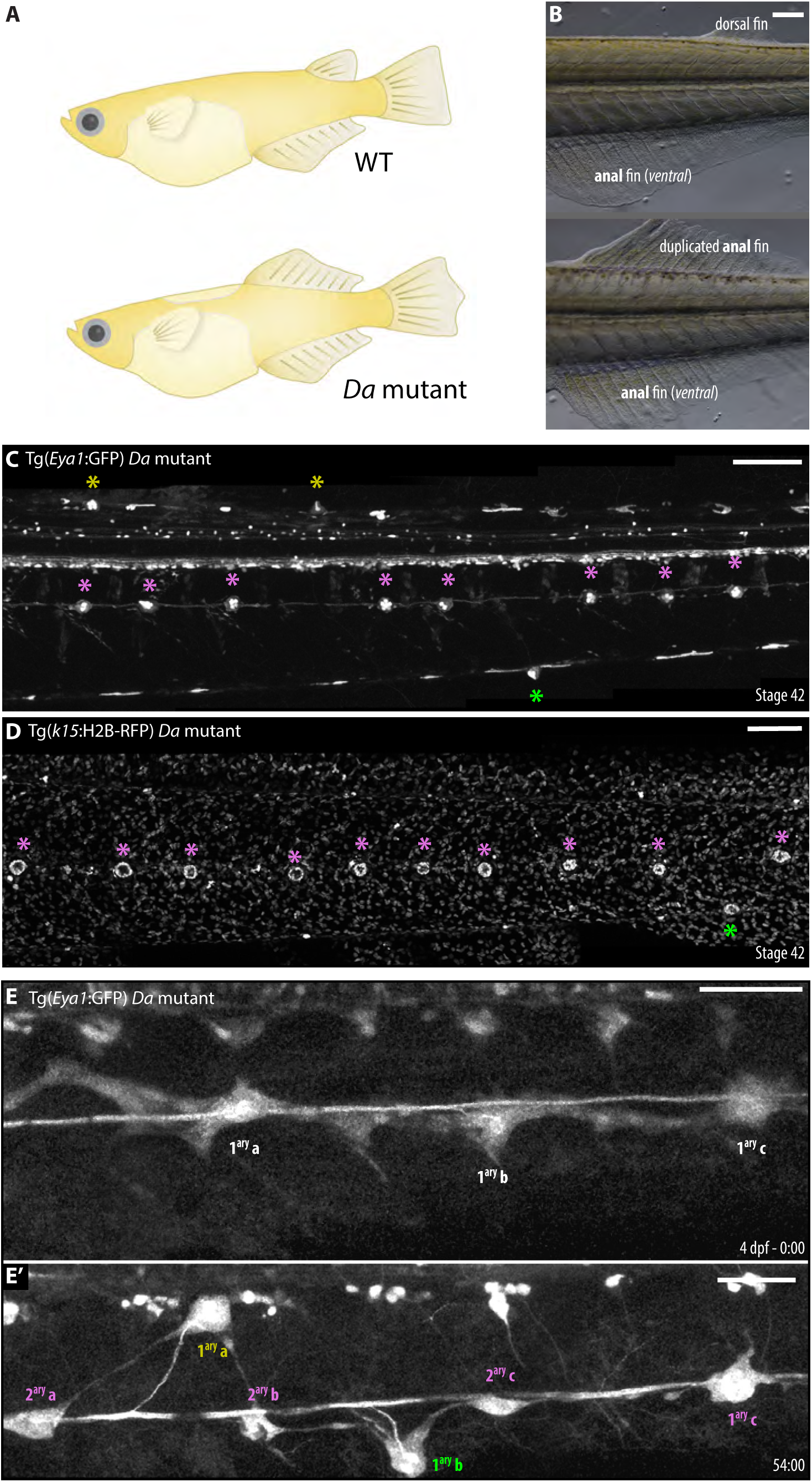
Location of primary neuromasts in *Da* mutants reveal autonomous organ positioning in the pLL system. (**A**) Scheme representing main differences in body shapes between adult *wt* and *Da* mutant fish. Notice the duplication of the anal fin dorsally in the *Da* mutant. (**B**) Bright-field view of wt and Da medaka larvae showing dorsal and anal fin positions. Scale bar = 100 microns. (**C**) Tg(*Eya1*:GFP) *Da* mutant shows neuromasts locating to both ventral and duplicated ventral side (green and yellow asterisks, respectively) while many neuromasts remain stuck in the midline (magenta asterisks). Scale bar = 100 microns. (**D**) A Tg(*krt15*:H2B-RFP) *Da* mutant shows additional organs locating to the midline (magenta asterisks). Scale bar = 100 microns. C & D panels, N > 50 *Da* larvae. (**E-E’**) Snapshots of a 4D-live SPIM imaging on 4 dpf Tg(*Eya1*:GFP) *Da* mutant during pLL organ deposition. Notice primary organs moving to the induced ventral side (yellow label), the regular ventral side (green label) and an organ that remains stuck in the midline (magenta label). Scale bar = 50 microns. Time in hours. N=5 larvae.

Interestingly, in all cases observed in the *Da* mutant, a secondary organ was formed in between two primaries regardless of their position (ventral-ventral, ventral-dorsal, dorsal-dorsal and dorsal-ventral) (N= 5 embryos) (Supplementary Movie 1). These results point to the fact that intrinsic properties of the primordium (one secondary organ in between two primaries, secondary organ positioning) and extrinsic properties (positioning of primary organs) together govern the construction of the pLL pattern in medaka. Also, they highlight the plasticity of this neural system to respond differentially to local changes in its environment on an organ-by-organ level creating a novel pLL pattern in the process.

### *K15* mutants show a perturbed epithelium and pLL patterning defects

Intrigued by the possibility that modulating the immediate environment of the primordium might lead to a modulation of pLL patterns, we decided to disrupt epithelial integrity by targeting *keratin 15 (krt15)* using Crispr/Cas9 (see M&M). Both F0 crispants and stable mutants (Supplementary Figure 1) showed epithelial extrusion (Figure 4A,B white arrows), perturbations in the structural integrity and cellular packing in the supra-basal epithelial layer (Figure 4 C-D, N=>100 cells segmented in *wt* and *krt15* mutants, N= 5 control & 15 *krt15* mutants larvae), lesions in the basal epithelium (Figure 4 F-G’ N>10 embryos) and epithelial cell death (Figure 4 G-G’ magenta arrows N=5 embryos) as revealed by nuclear rounding and cell membrane shrinkage, all in line with known Krt15 functions in epithelial cells (Bose et al., 2013; Chamcheu et al., 2011; Giroux et al., 2017; Giroux et al., 2018; Haines and Lane, 2012; Liu et al., 2003; Peters et al., 2001). In addition, *krt15* crispants and mutants showed a perturbed lateral line pattern, with primary organs locating to the midline (Figure 4 E-G’). While primary and secondary organs are arranged in an alternating pattern in *wild types* (N=24/24 pLLs), we have observed primary organs retained in the midline of *krt15* homozygous mutants (N=25/26 *krt15* pLLs)(see M&M).

**Figure 4.**
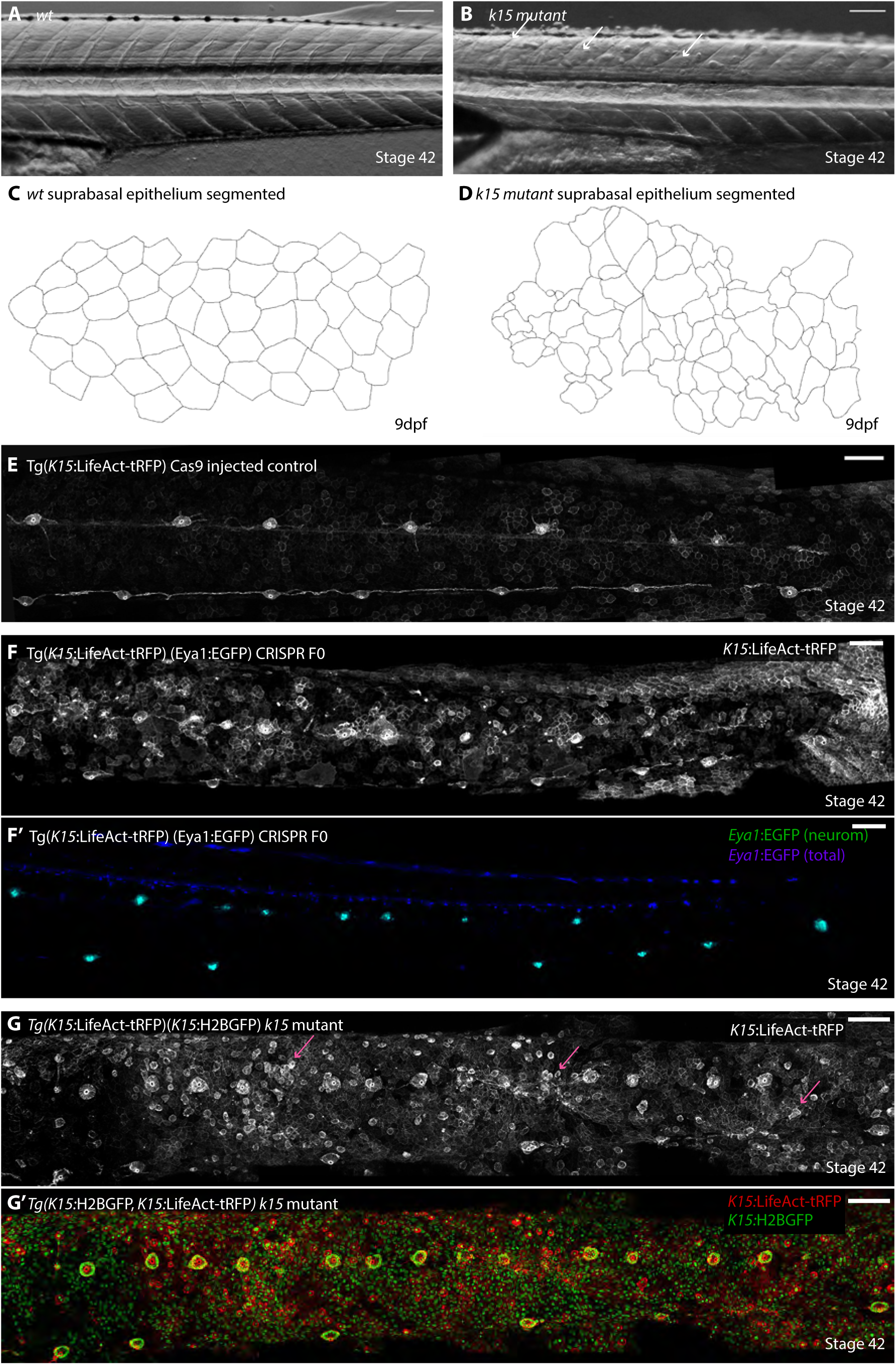
*Krt15*^-/-^ mutants show epithelial and pLL patterning defects. (**A**) Bright-field imaging of wt stage 42 larvae, notice normal epithelium. (**B**) Bright-field imaging on *krt15* mutant stage 42 larvae, notice the presence of extruding cells from the epithelial cell layer (white arrows) (C) Manual segmentation of *wt* suprabasal epithelial layer in 9 dpf embryos, notice the regular packing and conserved shape of epithelial cells. (**D**) Manual segmentation of *krt15* mutant suprabasal epithelial layer reveals massive disorganisation, loss of structural packing and loss of uniform cell size N >100 cells segmented in *wt* and *krt15* mutants, N= 5 control & 15 *krt15* mutant larvae. (**E**) Control Cas9 injected Tg(*K15*:LifeAct-tRFP) with alternating pLL pattern. (**F, F’**) Double Tg(*K15*:LifeAct-tRFP)(Eya1:EGFP) crispant injected with Cas9 + gRNA1,2 against *krt15* locus. (F) Notice epithelial disorganization and lesions, disturbed pLL patterns with many organs in a row at the midline. (F’) Same larvae showing *Eya1*:EGFP, whole expression is in blue and neuromasts are displayed in green to highlight pLL patterning defects. N>100 injected embryos, 40% showing a phenotype in pLL patterning and skin lesions. (**G-G’**) Double Tg(*K15*:H2B-GFP)(*K15*:LifeAct-tRFP) *krt15* mutant recapitulates phenotypes observed in the injected (F0) generation. Notice the presence of epithelial lesions (F, magenta arrows) and the heavily perturbed pLL pattern (F’) where many primary neuromasts are located at the midline. Scale bar = 100 microns.

**Figure 5.**
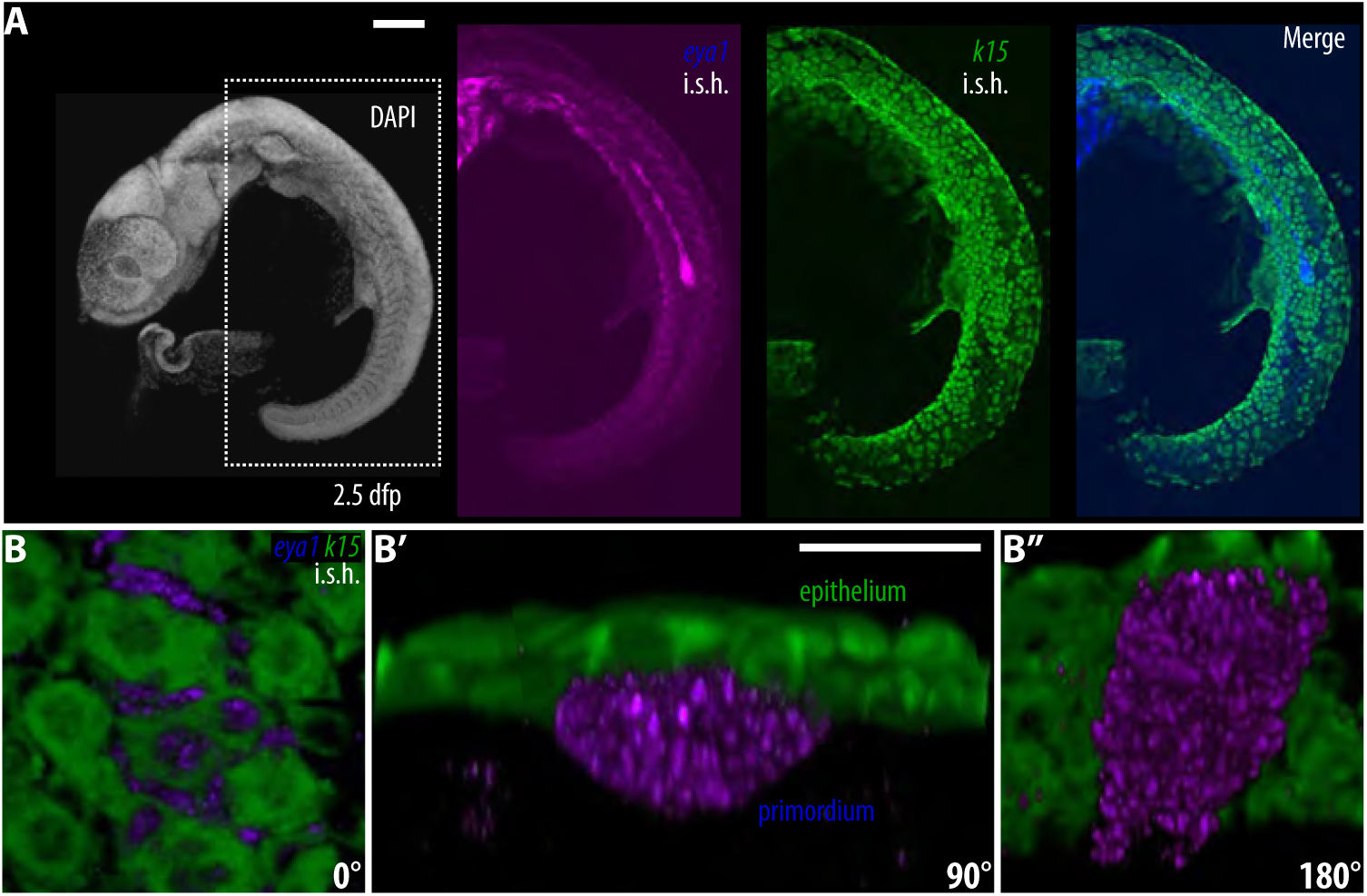
Perturbed *krt15* mutant epithelium causes pLL patterning defect. (**A**) Double *in situ* hybridization using *eya1* and *krt15* antisense probes during the primordium migration stage. *eya1* is strongly expressed in the primordium while *krt15* is mainly in the epithelium with no detectable expression in the primordium. Scale Bar = 100 microns (**B-B’’**) 3D reconstruction on the primordium of a sample treated as in (A). 0°, 90°, and 180° rotations to show differential expression of *eya1* in the primordium and *krt15* in the epithelium. Scale bar = 20 microns.

By tracing the positioning of the nerve connections in *krt15* mutants it seemed plausible that primary organs are initially specified correctly and move ventrally but at a later point revert their position and locate to the midline. To test this hypothesis directly and to reveal the basis for the pLL phenotype in *krt15* mutants we decided to rely on a 4D-live imaging approach. This revealed that primordium migration is significantly slower and stuttered in *krt15* mutants as compared to wt and *Da* mutants (4,71 +/- 2,38 µm/h in *krt15* mutants, N = 3 embryos, 17,95 +/- 3,69 µm/h in *wild types*, N = 5 embryos and 18,28 +/- 4,71 µm/h in *Da* mutants, N = 5 embryos) (Supplementary Movies 3 & 4, N = 3 *krt15* mutant fish and 3 *wt* fish), coinciding with the presence of epithelial lesions that block the path of the primordium through the epithelium. Our live-imaging also revealed that a fraction of primary deposited organs starts moving ventrally but are heavily delayed or blocked from proceeding further, in some cases even reverting their direction to migrate towards the mid-line (Supplementary Figure 2). These results are in line with our nerve tracing experiments that show a reversion of the nerve connection from ventral to dorsal directions in *krt15* mutant embryos (Supplementary Figure 2). Intriguingly, one secondary organ forms in between two primary organs in all the *krt15* mutants, and these secondary neuromasts locate properly to the midline, strongly suggesting that both properties are indeed intrinsic features of the pLL system itself. While we have previously reported that neuromast numbers are variable between left and right pLLs within the same *wt* fish (Seleit et al., 2017a), both sides have an identical, alternating distribution between ventral primaries and midline secondaries. Interestingly, we have seen cases in which the distribution of neuromasts were different between left and right pLLs within Da or *krt15* mutant embryos (5/7 and 5/13 embryos, respectively), typically showing one side with more mislocated primary organs that the other. Both the asymmetries in organ distribution between left and right pLLs within a fish, and the fact that some but not all primary organs are mislocated along a given pLL, highlight the degree of phenotypic plasticity in the system and suggest once more that subtle local differences in immediate micro-environment can have a large impact on resulting patterns.

### Epithelial cell integrity modulates pLL pattern in medaka

A double *in situ* hybridisation using specific anti-sense probes for *eya1* and *krt15* showed that *krt15* mRNA is not detectable in the primordium tissue during migration (Figure5A-B’’ N=3), indicating that the pLL patterning defect observed in the *krt15* mutant might indeed be extrinsic to the primordium. To directly test whether the patterning defects in the *krt15* mutant arise intrinsically due to defects in the primordial tissue or extrinsically from the local environment the primordium travels through, we decided to perform blastula stage transplantation assays from *krt15* mutants into wt and vice-versa, since we have previously shown that the neuromasts and the surrounding epithelium have two independent embryonic origins (Seleit et al., 2017b). Our results show that the presence of a large majority of *krt15* mutant cells within a primary neuromast does not lead to any patterning defects (Figure 6A-B’’ N=6 chimeric embryos n=12 mosaic organs), since neuromasts correctly position to the ventral side and the pLL pattern observed is equivalent to the wild-type. On the other hand, the presence of a vast majority of wild type cells in the neuromast in a *krt15* mutant background fails to rescue the patterning defect (Figure 6C-D, N= 3 chimeric fish and n=6 mosaic organs). Along the same lines, we obtained one transplanted fish in which *wild type* cells contributed to the epithelium of the right side of the tail in an otherwise *krt15* mutant background. In that larva, the left pLL displayed the typical aberrant organ distribution reported for *krt15*, with most neuromasts located along the midline (Figure 6E-E’’). On the right side, however, the presence of a *wild type* epithelium rescued the alternating distribution of neuromasts, where primary organs locate ventrally and secondary neuromasts locate at the middle (Figure 6F-F’’). These data combined with previous results strongly argue that the observed pLL patterning defect arises from the local environment the primordium and primary deposited organs travel through, namely the epithelium.

**Figure 6.**
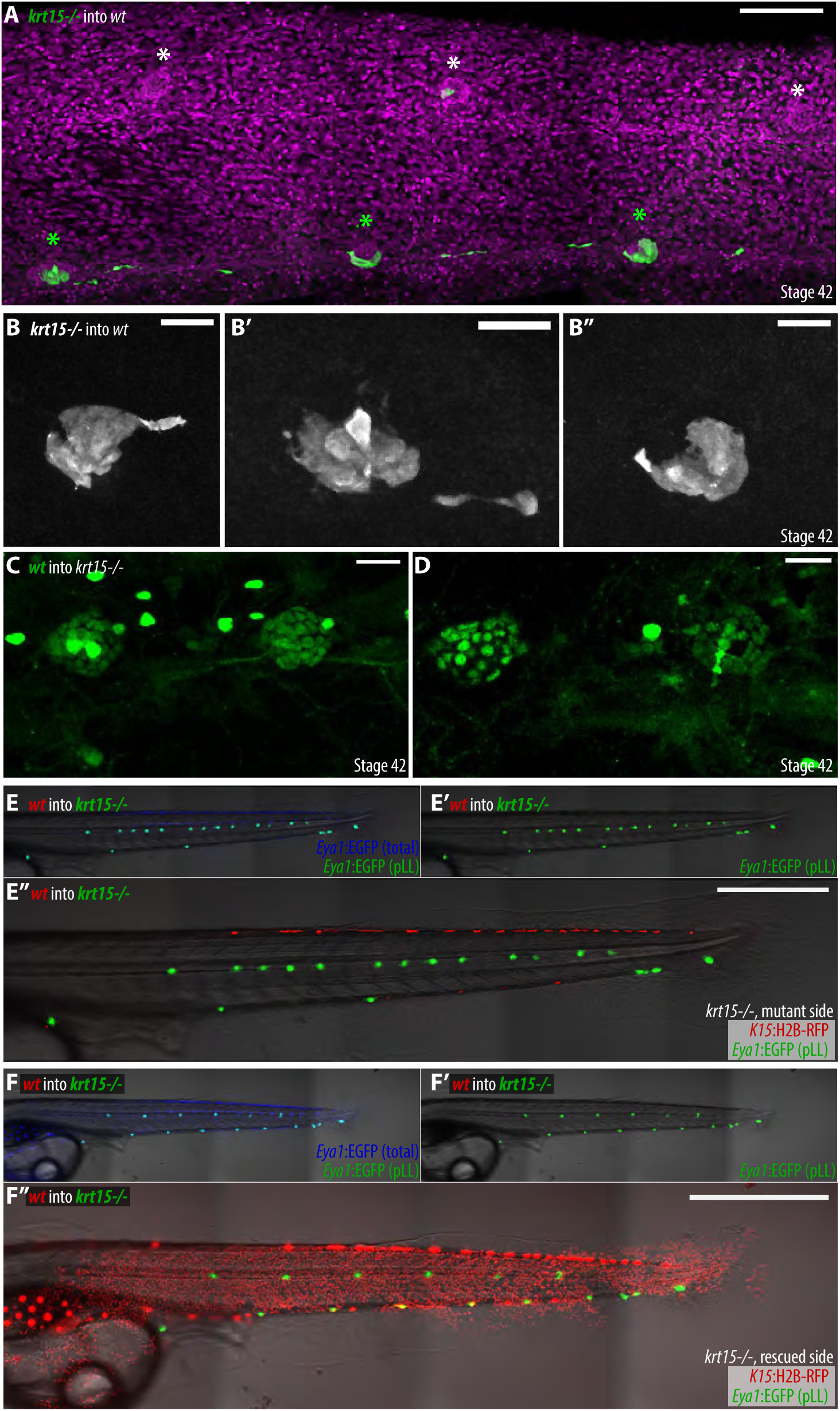
Perturbed *krt15* mutant epithelium causes pLL patterning defect. (**A-B’’**) Anti-EGFP antibody staining on chimeric larvae where Tg(*Eya1*:GFP) *krt15* mutant cells were transplanted into unlabelled *wild-type* background. (A) Organs with majority of *krt15* mutant cells position correctly to the ventral side. N = 6 transplanted embryos, N=12 mosaic organs. Scale bar = 100 microns. (B-B’’) Close up on ventral mosaic organs showing majority of krt15 mutant cells. Scale bar = 20 microns. (**C-D**) In vivo imaging of chimeric larvae where Gaudi^LoxPOUT^ donor cells (ubiquitously expressing H2B-EGFP) were transplanted into unlabelled *krt15* mutants. Organs containing mostly *wild type* cells remain at the midline. N = 3 transplanted fish and n = 6 neuromasts. Scale bar = 20 microns. (**E-F’’**) A chimeric *krt15* mutant larvae that displays wild type cells along the right trunk rescues the alternating distribution of neuromasts. EGFP expression in Tg(*Eya1*:EGFP) is displayed in blue (E, F), while EGFP positive neuromasts are represented in green (a subgroup of EGFP positive cells)(E’, E’’, F’, F’’). (F’’) The *krt15* mutant side shows the aberrant neuromast location. (F’’) *krt15* mutant neuromasts arrange in an alternating manner when surrounded by wild type epithelial cells (red). Scale bar in E’’, F’’ = 1 mm.

Complementing the *krt15* genetic perturbation, we decided to mechanically interfere with epithelial cell integrity, to address whether the disruption of epithelial integrity was the driving force behind the aberrant neuromast distribution in *krt15* mutants. Using 2-photon laser ablation we targeted *krt15* positive epithelial cells directly below deposited primary organs during primordium migration in 3-4 dpf *Krt15*:mYFP embryos (Figure 7A), leaving both the primordium and deposited organs uninjured and intact. Iterative imaging 48 hours post ablation revealed that mechanically induced lesions interfered locally with correct ventral migration of primary organs, and therefore recapitulated our results from the *krt15* mutant (Figure 7B-C’ N=5 ablated embryos and N=3 non-ablated controls). Notably, neuromasts anterior and posterior to the injury site, not exposed to a mechanical perturbation in their immediate surrounding, positioned correctly along the body axis (Figure 7C). Overall, our results indicate that the pLL pattern is easily modulated by changes in tissues external to the primordium (*i.e* the epithelium), suggesting that this adaptability can be exploited evolutionarily to match a somatotopic sensory system to variable body shapes.

**Figure 7.**
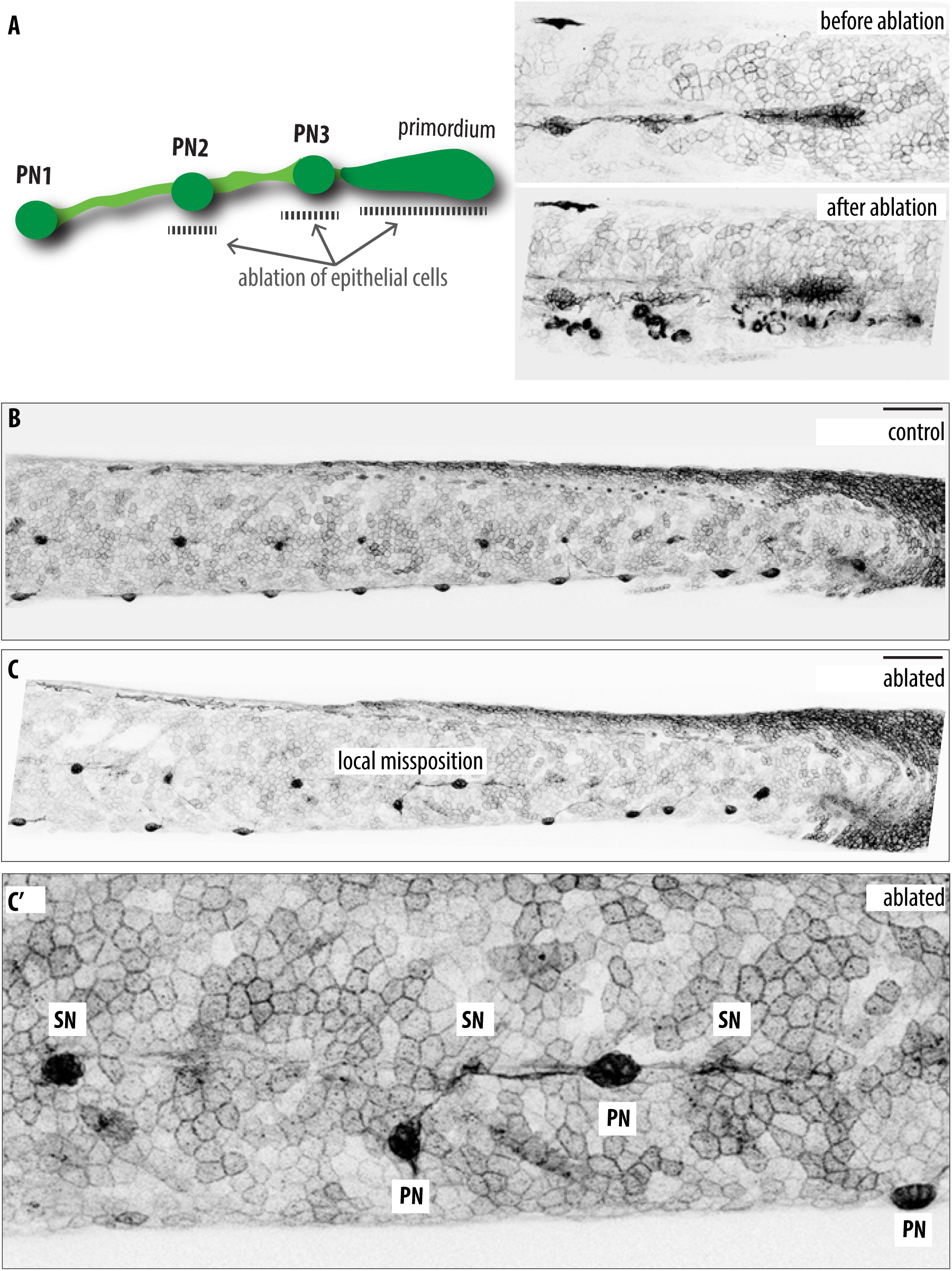
Local mechanical perturbation of epithelium leads to pLL patterning defects. (**A**) Schematic diagram of 2-photon ablation on Tg(*K15*:mYFP), 3dfp (left). Tg(*K15*:mYFP) embryos showing primordium migration and primary deposited organs before (upper right) and immediately after (bottom rigth) 2-photon laser ablation of epithelial cells below primary deposited organs. (**B**) 5 dpf control, unablated Tg(*K15*:mYFP) showing alternating neuromast pattern with primary organs locating ventrally. (**C**) 48 hours post epithelial cell ablation, two primary organs fail to migrate ventrally at the site of injury, while organs anterior and posterior to the injury site locate normally. (**C**’) Close up view of primary organs that failed to migrate ventrally in (C). N=3 Ablated fish and 3 controls. Scale bar = 100 microns.

## Discussion

How biological patterns are built and how they evolve is a central question in developmental biology. Recent work has argued for cis-regulatory element refurbishment as a means for evolutionary tinkering (Chan et al., 2010; Indjeian et al., 2016; Jones et al., 2012; McLean et al., 2011; O’Brown et al., 2015; Prud’homme et al., 2006). Yet it is still unclear how reproducibility within one species is balanced with the requirements of evolution of forms and patterns between species. In this work we show that for a peripheral nervous system in vertebrates, the strategy relies on balancing the strict internal properties of the system with a plastic response to the immediate microenvironment executed autonomously on an organ-by-organ basis. The self-organizing and hierarchical nature of the process ensures that small changes in conditions can lead to large variations in patterns without affecting organ morphogenesis. Our data support the hypothesis that these changes do not need to occur within the pLL founding tissue *per se* and do not need to involve any regulatory changes within the genetic landscape controlling primordium migration. The unpredictability of individual organ positioning in *Da* and *krt15* mutants points towards the fact that the posterior lateral line system is phenotypically plastic. This low developmental buffering capacity (Waddington, 1959; Waddington, 1968, 1969) for the pLL would ensure it is able to evolve rapidly in response to changes in other tissues or overall morphology of teleost fish. That these changes happen during embryogenesis emphasises the already well-established paradigm of development as the playing field of evolution (Carroll, 2005). Our results are surprising since they show that a new pattern can be obtained without changing any internal or genetic properties of the cells executing organogenesis. Instead, the system has the capacity to sense, react and respond to changes in its surroundings in an organ autonomous manner. By probing differential organ positioning the pLL system is well suited for a rapid evolutionary diversification; indeed, by looking at different embryonic pLL patterns in a number of species we were able to see evidence of this plasticity at play (Figure 8).

**Figure 8.**
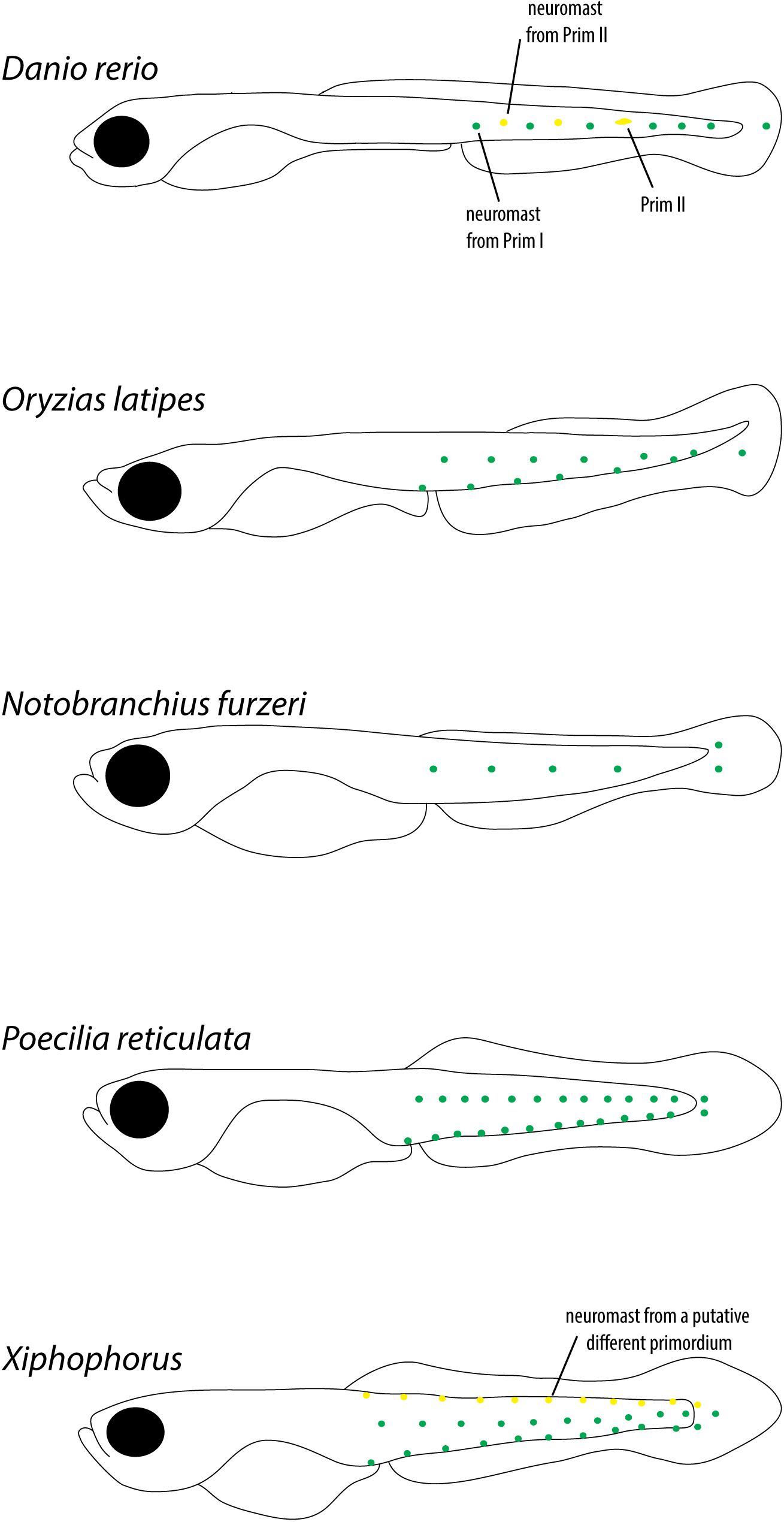
Diversity of embryonic pLL pattern and organ number in different teleosts. The diversity of pLL patterns at the end of embryogenesis in a variety of teleosts. Scheme showing the distribution and approximated organ numbers in a variety of teleost fish that were stained using either DAPI or DiAsp and imaged. N = 4 for each species, but zebrafish and medaka N > 20. Fish embryos are not drawn to scale.

The main function of the lateral line in teleost fish is to sense the direction of water flow and relay this information back to the brain (Ghysen and Dambly-Chaudière, 2007). Critical to the fulfillment of this function is proper, species-specific numbers and distribution of neuromasts. Previous work in zebrafish reported that a fixed number of organs is deposited by the primordium (Ghysen et al., 2010; Gompel et al., 2001; Sarrazin et al., 2010) but that the number of subsequent, inter-calary organs relies on external cues (Ghysen et al., 2010; Nuñez et al., 2009; Sapède et al., 2002). Specifically, it has been shown that the loss of the pLL nerve and associated glia in a variety of mutants leads to supernumerary neuromasts (Grant et al., 2005; López-Schier and Hudspeth, 2005; Lush and Piotrowski, 2014; Whitfield, 2005), the source of which is an over-proliferation of latent inter-neuromast cells (Grant et al., 2005; Lush and Piotrowski, 2014)). While there is an obvious effect on organ numbers, the positioning of these extra organs in zebrafish remain largely equivalent to the wild-type, where all neuromasts are arranged along the horizontal myoseptum at the end of embryogenesis (Grant et al., 2005; López-Schier and Hudspeth, 2005). Recent work has shown that Erbb signaling emanating from glial cells contributes to maintaining organ numbers in check (Lush and Piotrowski, 2014). Our results in medaka challenge the generality of these findings in other teleosts. First, we clearly show that the pLL nerve is dispensable for proper organ numbers and positioning. Second, we show that a normal pLL pattern and organ numbers are maintained in the absence of S*ox10+* associated glia. It is therefore plausible that different mechanisms exist that control pLL pattern and organ numbers in different teleosts. However, we and others have reported that *ngn1* mutants and crispants show an increase in organ numbers in the pLL of both medaka and zebrafish (López-Schier and Hudspeth, 2005; Seleit et al., 2017b). Since *ngn1* acts developmentally earlier during placodal specification (Andermann et al., 2002; López-Schier and Hudspeth, 2005) these defects might therefore not solely pertain to the presence or absence of glial cells. We have previously reported that the posterior lateral line primordium present in medaka resembles zebrafish PrimII in terms of the overall pattern generated (Nuñez et al., 2009; Seleit et al., 2017a). Intriguingly none of the previously described mutants leading to supernumerary neuromast numbers in zebrafish were reported to have an effect on PrimII derived neuromasts (Grant et al., 2005; López-Schier and Hudspeth, 2005). This raises the possibility that neither PrimII in zebrafish nor the medaka pLL primordium are affected by loss of the pLL nerve or associated glia. Related to the nerve ablation experiments, two additional observations are worth discussing. First, it has been reported previously in zebrafish that peripheral nerves act instructively during post-embryonic neuromast formation (Wada et al., 2013a), since organs will not form without a preceding nerve outgrowth. In the embryonic pLL development in medaka, however, the nerve is entirely dispensable for organ formation, organ numbers and patterns, highlighting that even within similar systems, the hierarchical logic governing the sequence of events required for their construction can be markedly different. And secondly, an unexpected outcome of the nerve ablation experiments is the observation that medaka pLL nerve is not able to re-grow after injury, a situation that differs from what is reported in zebrafish (Gilmour et al., 2002). This deficiency in regenerative potential in medaka has been reported for a number of other organs (Ito et al., 2014; Lai et al., 2017; Lust and Wittbrodt, 2018) and the peripheral nervous system can now be added to the growing list of regeneration-deficient components. The molecular underpinnings of this fundamental difference in regenerative capacities between different teleosts is only beginning to be explored (Lai et al., 2017; Lust and Wittbrodt, 2018).

A kaleidoscopic diversity of body plans exists in teleost fish (Coombs et al., 2014; Coombs et al., 1988). This raises immediate problems for the set-up of a peripheral nervous system component like the neuromasts of the lateral line during development (Ghysen and Dambly-Chaudiere, 2016). On the one hand the system must be tunable and easily adjustable to fit the diverse body plans of the different species, and on the other it must still be reproducible within and between members of the same species. We show that the system accommodates these needs by balancing fixed intrinsic properties (reproducibility) with an ease of response to small changes in extrinsic cues (adaptability). In medaka, neuromasts invariably alternate between the midline and the ventral line irrespective of the total number of organs that the posterior lateral line harbours (Seleit et al., 2017a). By challenging the medaka lateral system to different contexts we were able to gain new insights into the operational logic behind pLL pattern construction. This was done by using a medaka mutant with an unconventional body plan, a mutant with a disturbed epithelial integrity, in addition to local mechanical perturbations to epithelial cells. And interestingly, we observe that even within the same *Da* or *krt15* genotype organ numbers and distribution can be decoupled between left and right pLLs of the same fish. This strongly suggests that local interactions and other non-genetic causes might contribute to the phenotypic plasticity of this developmental morphogenetic process. The *Da* mutant challenges the system by the presence of two ventral sides on the tail, revealing the autonomous nature of organ positioning, *i.e.* primary neuromasts locate to the ventral, dorsal (induced ventral) or midline independently of one another. Organ positioning is therefore resolved in a neuromast neighbor-independent manner arguing against the presence of a system level control on the process. Our results point to a high potential of plasticity in the system but within defined and strict boundaries ensuring a reproducible outcome. Specifically, it seems the system balances intrinsic fixed properties with extrinsic malleability and adjustability. We report that the generation of a single secondary organ in between two primaries seems to be an intrinsic property of the medaka pLL primordium, observed in the *Da* mutant, the *krt15* mutant and in the ablation of the posterior lateral line nerve. Extrinsically, the immediate environment the primordium and primary deposited organs travel through directly influences the final position of the organs and therefore the overall pLL pattern at the end of embryogenesis. A sequential order emanates from this logic, where first, local extrinsic parameters modulate primary organ positioning, second, neuromast migration dictates the innervation and therefore neuronal network (Haas and Gilmour, 2006) and third, the final position of the organ induces in situ the formation of a life-long niche for neuromast stem cells during organ maturation (Seleit et al., 2017b). The same hierarchical organisation is well reported in different induction paradigms, like the induction of the lens by the neural retina in vertebrates (Cvekl and Ashery-Padan, 2014) or the generation of a new neural plate by the Henses’ node in chicken (Storey et al., 1992) (Anderson and Stern, 2016).

We have previously reported that *krt15* is expressed in the basal (dividing) epithelial cell layer of medaka in addition to the long-term stem cells within mature neuromasts (Seleit et al., 2017b), in line with its known role as an epithelial stem cell marker in a variety of tissues (Bose et al., 2013; Giroux et al., 2017; Giroux et al., 2018; Liu et al., 2003). Our newly generated *krt15* mutant shows epithelial lesions and the loss of epithelial cell integrity in addition to epithelial cell death and extrusion, phenotypes consistent with its functional roles in the epithelium of other model organisms (Chamcheu et al., 2011; Haines and Lane, 2012; Peters et al., 2001). Interestingly we also report a strong phenotype in the pLL pattern of medaka and by transplantation assays and mechanical disruption of the skin we trace its origin to the perturbed mutant epithelium. Recent results in zebrafish revealed that the epithelium is necessary for the migration of the primordium, since skin removal results in halting primordium migration, which will resume only after re-growth of the epithelial cells (Nogare et al., 2019). Consistent with this, we observe phenotypes in primordium migration, *i.e.* stuttering, in the medaka *krt15* mutants in response to epithelial cell lesions. In addition to this our results strongly suggest that the degree of structural integrity of the tissue the primordium and deposited primary organs travel through can directly modulate pLL patterns.

The results we report along the manuscript suggest that pLL pattern construction is a system with a low developmental buffering capacity. The concept of developmental buffering was first described by Conrad Waddington to explain the canalization of developmental processes i.e. the arrival at the same reproducible outcome despite small genetic and non-genetic differences in starting conditions (Waddington, 1941, 1959; Waddington, 1968, 1969). However, Waddington also recognised that different developmental modules must have different buffering capacities depending on need (Waddington, 1959). In the case of the pLL it would be evolutionarily beneficial to operate under a low developmental buffering capacity as it would make the system highly susceptible to small changes in conditions. It is tempting to speculate that the high plasticity of the system could be exploited evolutionarily to generate diverse patterns of lateral lines in the wild. Combining examples from the literature and our own cross-species observations confirms that embryonic pLL patterns are variable among different teleost species (Figure 8), highlighting once more the degree of plasticity and fast evolvability of this developmental module.

## Acknowledgements

We would like to thank Steffen Lemke, and the entire Centanin and Lemke lab members for critical input on the manuscript and for fruitful discussions. We would like to thank Jochen Wittbrodt for the generous support and access to equipment, Hiro Takeda for *Da* mutants, Tatjana Piotrowski for the *Sox10* promoter, Manfred Schartl for providing fixed samples of *a* number of teleost species used in this study, NBRP Medaka for materials and E. Leist, A. Sarraceno and M. Majewski for assistance regarding fish maintenance. This work has been funded by the Deutsche Forschungsgemeinschaft (German Research Foundation, DFG) via the Collaborative Research Centre SFB873.

## Supplementary Figure Legends

**Supplementary Figure 1.**
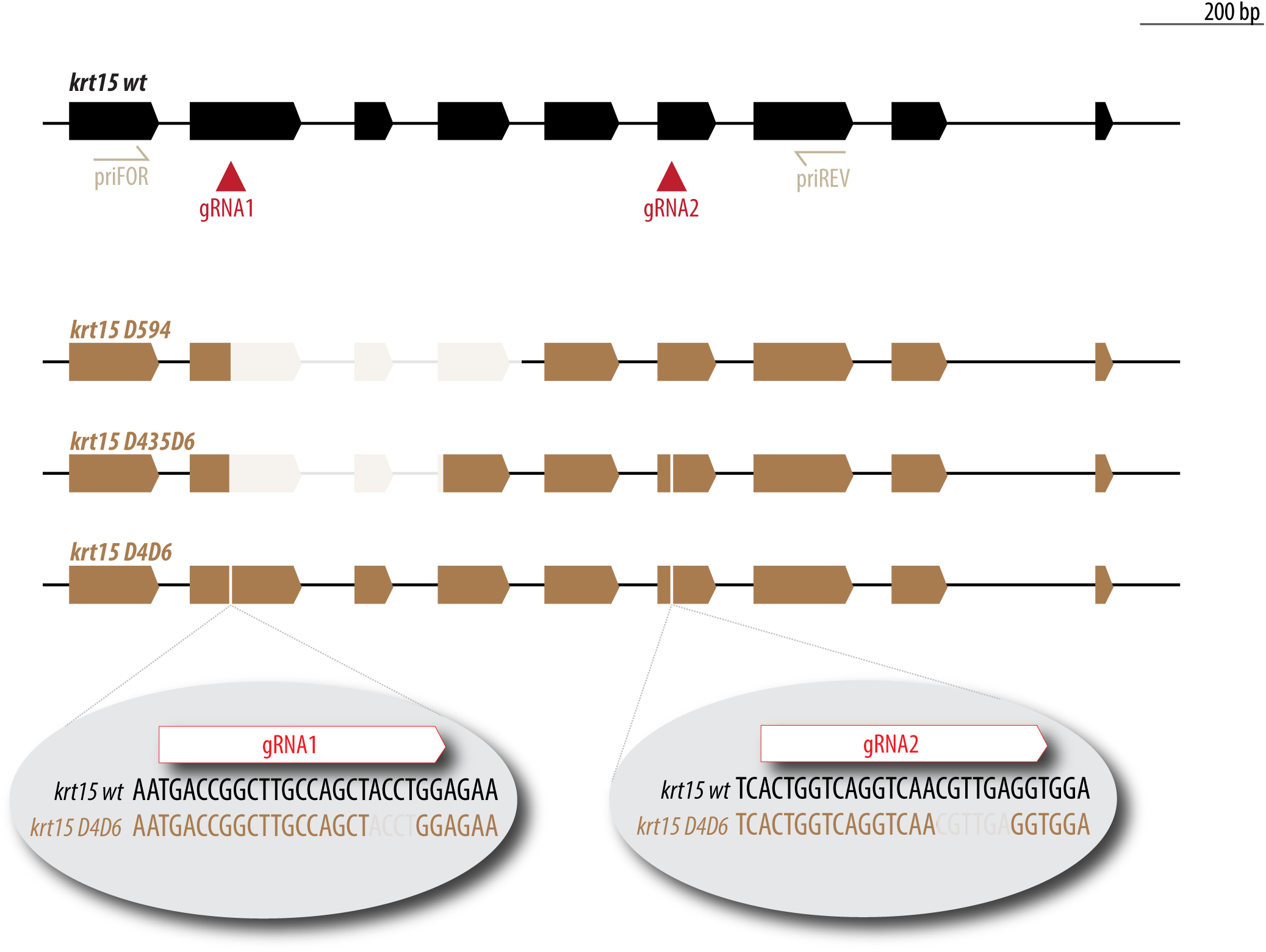
We have targeted the medaka *krt15* locus (top) with two different gRNAs (red triangles). Among the mutant alleles obtained, we isolated and grew stable mutant lines for the following alleles that are displayed in the figure: *krt15*^D594^, *krt15*^D435D6^, *krt15*^D4D6^. Major deletions are represented by a shadow box (*krt15*^D594^, *krt15*^D435D6^), while for minor deletions of 4 - 6 bp (sgRNA2 site for *krt15*^D435D6^, and both gRNA sites for *krt15*^D4D6^) we present the DNA sequence in the altered region (bottom).

**Supplementary Figure 2.**
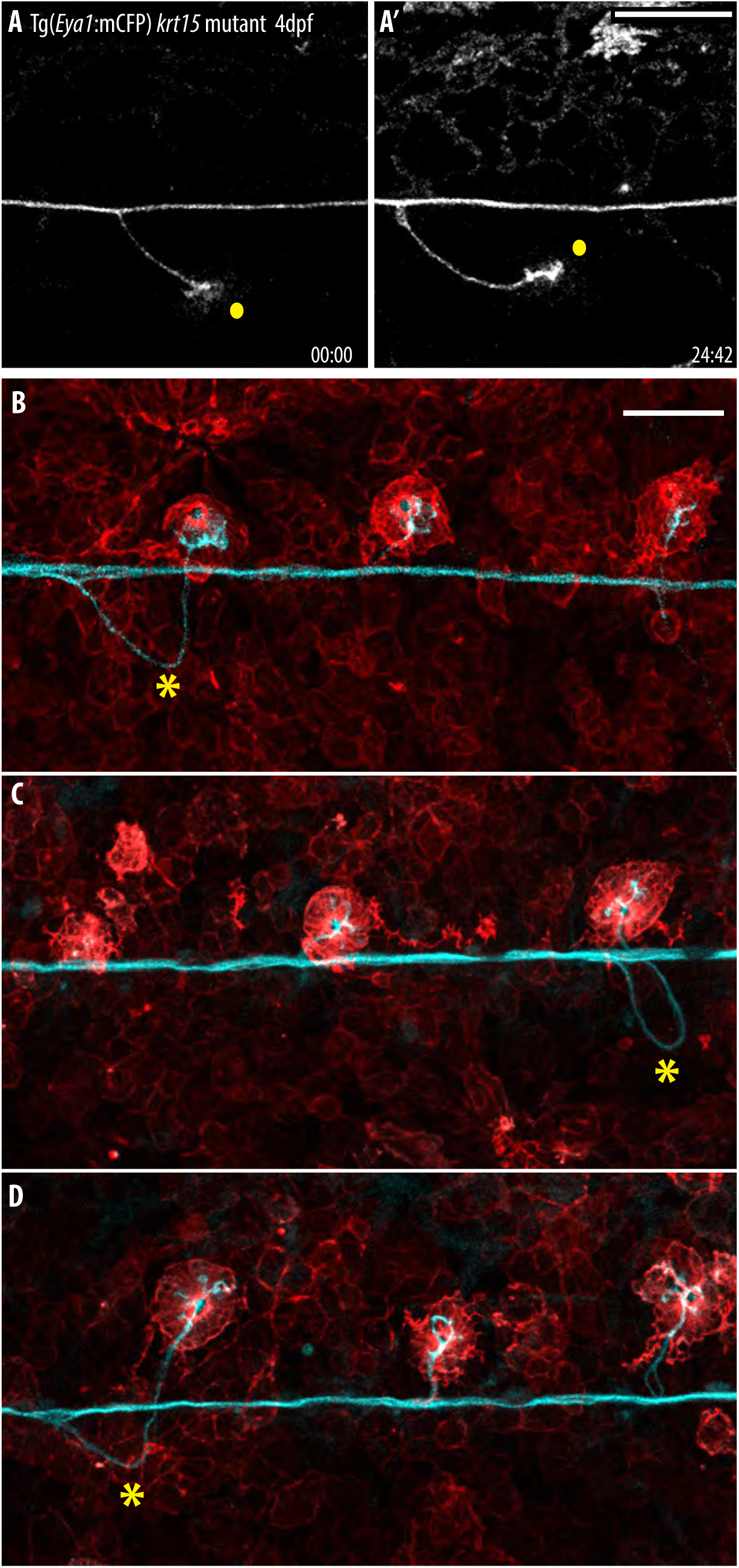
(A-A’) Stacks from time-lapse confocal imaging of Tg(*Eya1*:mCFP) *krt15* mutant during primary organ ventral displacement. Notice that in cases, primary neuromasts start ventral displacement (yellow dot, A) as in the wild type, but revert later to the midline (yellow dot, A’). N= 6 fish. Time in hours. Scale bar=50 microns. (B-D) Confocal image of Tg(*Eya1*:mCFP)(*K15*:LifeAct-tRFP) *krt15* mutant larvae. Primary organs are located in the midline, and their nerve path reveals a prior ventral displacement.

## Supplementary Movie Legends

**Supplementary Movie 1. Uncoordinated primary neuromast migration in *Da* mutants.**

Time-lapse SPIM imaging of 3 dpf Tg(*Eya1*:GFP) *Da* mutant during primordium migration reveals that primary organs resolve positioning as autonomous units. Notice that the pLL nerve will migrate together with the neuromasts regardless of migration direction. A secondary organ will form between 2 primary organs irrespective of their migratory direction. Scale bar= 30 microns. Time in hours.

**Supplementary Movie 2. Primary organ movement in *Da* mutant**

Time-lapse SPIM imaging of 3-4 dpf Tg(*Eya1*:GFP) *Da* mutant during primary organ deposition. Zoom-in on primary organ movement: notice the change of position relative to the horizontal myoseptum, initially the organ moves ventrally, subsequently it moves in the dorsal direction before eventually moving back towards the horizontal myoseptum where it will locate. Scale bar= 30 microns. Time in hours.

**Supplementary Movie 3. Primordium stalling and primary organ positioning defects in *krt15* mutant embryo**

Time-lapse confocal imaging of 3 dpf Tg(*Eya1*:mCFP) *krt15* mutant during primordium migration. Notice that the primordium stalls and stops migration before resuming its movement. In addition, primary organs start ventral displacement in the mutant as revealed by the pLL nerve connection, however their normal ventral movement is heavily perturbed. N=3 *krt15* mutants.

**Supplementary Movie 4. Primordium stalling and epithelial lessions in *krt15* mutant embryo**

Time-lapse SPIM imaging of 3 dpf Tg(*Eya1*:EGFP)(*K15*:H2B-EGFP) (k15-LifeAct-tRFP) *krt15* mutant during primordium migration. Notice that the primordium momentarily halts migration before resuming its movement. The nuclear and cellular membrane labelling in the epithelium helps visualising lesions in the skin and apoptotic nuclei forming around primary organs. N=2 *krt15* mutants.

## Materials and Methods

### Animal ethics and strains used

Medaka were maintained as closed stocks at the Centre of Organismal Studies of the University of Heidelberg (Tierschutzgesetz §11, Abs. 1, Nr. 1). Medaka and zebrafish husbandry and experiments were performed according to local animal welfare standards (Tierschutzgesetz 111, Abs. 1, Nr. 1, Haltungserlaubnis) and in accordance with European Union animal welfare guidelines. The fish facility is under the supervision of the local representative of the animal welfare agency. The fish colony was maintained under standard recirculating aquaculture conditions, 14 hours of light and 10 hours of darkness. The strains used in this study are: *Cab* (wild-type population), *Da* mutant (Ohtsuka et al., 2004).

### Transgenic lines

We have used the following transgenic lines: Tg(*Eya1*:mECFP), Tg(*Eya1*:EGFP), Tg(*K15*:H2B-EGFP), Tg(*K15*:H2B-RFP) (Seleit et al., 2017a; Seleit et al., 2017b), *Gaudi*^LoxPOUT^ (Centanin et al., 2014). The newly generated transgenic lines are:

#### Tg(*Sox10*:mCherry)

The plasmid *sox10*:mCherry (a gift from Thomas Look, Addgene plasmid # 98695) was injected in wild type or *Da*-/- medaka embryos at the one cell stage using I-SceI meganuclease protocols as previously described (Thermes et al., 2002).

#### Tg(*K15*:LifeAct-tRFP)

A LifeAct-tRFP peptide was clone under the previously published medaka *krt15* partial promoter (Seleit et al., 2017b) using Age1 and Not1 restriction enzymes. The resulting plasmid *K15*:LifeAct-tRFP was injected in medaka embryos at the one cell stage.

### Generation of *keratin 15* mutants

Two gRNAs targeting exons 2 and 5 of the *keratin 15* were designed using CCTop (Stemmer et al., 2015) and synthesised as previously described (Stemmer et al., 2015). The sequences of *krt15* gRNAs are UGACCGGCUUGCCAGCUACCUGG and CACUGGUCAGGUCAACGUUGAGG. The following oligos were used:

*sgRNA-1_F*: TAGGTGACCGGCTTGCCAGCTACC

*sgRNA1_R*: AAACGGTAGCTGGCAAGCCGGTCA

*sgRNA-2_F*: TAGGCACTGGTCAGGTCAACGTTG

*sgRNA-2_R*: AAACCAACGTTGACCTGACCAGTG

Annealed oligos were ligated into linearised DR274 plasmids (Addgene clone number: #42250) as described before (Seleit et al., 2017b). This was followed by *in vitro* RNA transcription using T7 MEGAshortscript Kit (Ambion) and cleaned-up using RNAeasy kit (Qiagen). The gRNAs were co-injected at 15 ng/μl each along with 150 ng/μl of CAS9 mRNA into Tg(*Eya1*:EGFP), Tg(*Eya1*:mCFP), Tg(*K15*:H2BGFP), Tg(*K15*:LifeActtRFP) one-cell-stage medaka embryos. To check for gRNA efficiency the following genotyping primers were used: *krt15* fwd, GGGACCAGAGTCTCTGTTTCC, *krt15* rev, TTGGTGTTCCATGTCGTTGC.

Individual adults of the F2 mutant lines were genotyped by PCR using *krt15 fwd* and *krt15 rev* primers the amplicons were analysed by Sanger sequencing. We identified several alleles that are described in Supplementary Figure 1. In the coming generations, pLL phenotypes were checked for the *krt15* mutation. *krt15*^D594^ / + adults were incrossed and their progeny sorted for primary neuromasts stucked in the midline. A genotyping PCR confirm that these fish were indeed *krt15*^D594^ mutants (15/16 *krt15*^D594^ homozygous, 1/16 *krt15*^D594^ heterozygous).

### Transplantation assay

Blastula stage transplantations in medaka were carried out as previously described (Rembold et al., 2006). Briefly 10-35 cells were transplanted from *Gaudi*^LoxPOUT^ (ubiquitously expressing H2B-GFP), or (*K15*:H2B-RFP) donors into *krt15* mutant hosts, and from *krt15* mutant donors (*Eya1*:GFP and *K15*:H2BGFP) into wt hosts. Transplanted embryos were kept in 1× ERM supplemented with antibiotics (penicillin-streptomycin solution from Sigma, P0781) and selected for fluorescent expression in the primordium. Positively transplanted fish with labelled donor cells in the neuromasts or in the vicinity of the neuromasts were analysed at stage 42 under a fluorescent binocular or under a confocal microscope.

### Live-imaging and sample preparation

For dechorionisation, embryos were rolled on commercially available sand-paper, washed in 1 X ERM, treated with hatching enzyme for 30-50 minutes at 28 °C, and washed abundantly in ERM to remove any residual enzyme. All live-imaging was done as previously described (Seleit et al., 2017a, b). Briefly, dechorionated embryos or hatchlings were anaesthetised in 1x ERM supplemented with Tricaine (Sigma-Aldrich, A5040) and then mounted in 0.6% low melting agarose on glass bottom dishes (MatTek corporation). Embryo screening was performed using either the Olympus MVX10 macrofluorescence binocular. Live-imaging was done using the MuVi-SPIM (Krzic et al., 2012) with two illumination objectives (10x Nikon Plan Fluorite Objective, 0.30 NA) and two detection objectives (16X Nikon CFI LWD Plan Fluorite Objective, 0.80 NA). Additional live-imaging was done using the confocal laser-scanning microscope Leica TCS SPE (40x oil objective) or Leica TCS SP5 II (10X dry, 40x dH2O objectives). Samples were placed in the Microscope Slide Temperature Controller from Biotronix to control temperature.

### Image and data analysis

All Image analysis was done using FIJI and ImageJ. Image stitching utilized 2D and 3D stitching plug-ins on ImageJ. All registration of imaging time-stamping, manual tracking, and manual segmentation of cells, was done using standard FIJI plug-ins.

### Stainings

#### Immunohistochemistry

IHC were performed on fixed medaka embryos as described before (Centanin et al., 2014). Primary antibodies used were Anti-eGFP Rabbit (Invitrogen, 1:500) and anti-tRFP Rabbit (Evrogen, 1:250). Secondary antibodies used were goat anti-Rabbit Alexa Fluor 488 (Invitrogen, 1:500), goat anti-Rabbit Alexa Fluor 546 (Invitrogen, 1:500). For DAPI staining a final concentration of 5 ug/l was used.

#### Double in situ Hybridisation

Samples were prepared as previously described (Seleit et al., 2017a). Hybridisation was performed with the *eya1* (DIG) and *krt15* (Fluo) probes in hybridisation mix overnight at 65°C. All following washing steps were done with TNT (0.1M Tris pH 7.5, 0.15M NaCl, 0.1% Tween20). The samples were blocked in 2% TNB (2% blocking reagent (Roche, REF 11 096 176 001) in TNT for 2-3 h at room temperature and incubated with an anti-Fluo-POD antibody (1/50, Sigma-Aldrich REF 11 426 346 910) in 2% TNB overnight at 4°C. On the next day after washing detection of the *krt15* probe was performed using the PerkinElmer TSATM-Plus Fluorescein System (NEL741B001KT). To stop all remaining POD enzyme activity the samples were incubated 20 minutes in 2% H2O2 in TNT. For visualization of the *eya1* probe the samples were incubated with an anti-Dig-POD antibody (1/50 in 2% TNB, Sigma-Aldrich REF 11 207 733 910) and DAPI (1/1000) overnight at 4°C. Detection of the *eya1* probe was performed using the PerkinElmer TSATM-Plus Cyanine 3 System (NEL744B001KT). The embryos were mounted in 0.6% low melting agarose in PTW and imaged at a confocal microscope.

#### DiAsp

Hair cells in neuromasts were visualised using the vital dye 4-Di-2-ASP (Sigma-Aldrich) as previously described (Sapède et al., 2002; Seleit et al., 2017a). Live samples were incubated for 5-10 min in a 5 mM DiAsp solution, washed in ERM and analysed using a fluorescent binocular.

### 2-photon laser ablation

For pLL nerve ablations a multi-photon laser coupled to a Leica TCS SP5 II microscope was used. Conditions varied across the replicates, the ‘point ablation’ function was utilised using 880 nm wavelength and a laser power between 30-50 % for a timeframe of 300-800 ms. A section of the pLL nerve was chosen and from 6-20 sequential points were ablated. Directly after injury the pLL nerve was imaged to check the efficiency of targeting the nerve. In other experiments, we targeted 5-15 EYFP positive epithelial cell membranes underneath the primordium and latest deposited neuromasts in Tg(*K15*:mEYFP) 3-4 dpf embryos during pLL pattern construction. Point ablation settings were used and laser power varied between 30-52 % with a time frame of 200-300 ms. Embryos were imaged before and after to check the success of targeting.

